# Loss of HOXB13 expression in neuroendocrine prostate cancer

**DOI:** 10.1101/2020.07.03.187302

**Authors:** Siyuan Cheng, Shu Yang, Yingli Shi, Runhua Shi, Yunshin Yeh, Xiuping Yu

## Abstract

HOXB13, the most posterior HOX B gene, is primarily expressed in the prostate. Using immunohistochemical staining, we evaluated the expression of HOXB13 in prostatic tissues. We found that HOXB13 was expressed in both benign prostatic tissues and prostate adenocarcinoma (AdPCa) but its expression was reduced or lost in neuroendocrine prostate cancer (NEPCa). Additionally, using RNAseq data of HOXs in cell lines derived from various tissue origins, we established HOX codes for these tissue types. We applied the HOX codes to PCa samples and found that the majority of AdPCa samples sustained the prostate-specific HOX code but the majority of NEPCa samples lost it. Our analysis also showed that the NEPCa samples did not correlate well with the HOX codes of any other tissue type. This indicates that NEPCa tumors lose prostate identity but do not gain a clear-cut new tissue identity. Finally, we treated PCa cells with all trans retinoic acid and found that the reduced/lost HOXB13 expression can be reverted, as can AR and AR targets. Taken together, our data indicate that HOXB13 expression is reduced or lost in NEPCa and a loss of prostate-specific HOX code in NEPCa represents a loss of prostatic identity.

## Introduction

Prostate cancer (PCa) is the most diagnosed cancer among American men. In 2020, 191,930 men in the US are predicted to be diagnosed with PCa and 33,330 men will die from this disease [1]. Neuroendocrine prostate cancer (NEPCa) is an aggressive type of PCa. After androgen deprivation therapy fails, about 30% PCa patients acquire the neuroendocrine (NE) phenotype [2]. Although the cell of origin of NEPCa has not been conclusively determined, accumulating evidence supports that NEPCa cells arise from the trans-differentiation of prostate adenocarcinoma[3]. Currently, there is no effective treatment for PCa with a prominent NE phenotype. Understanding how PCa progresses into NEPCa could have important diagnostic and treatment implications.

Homeobox-containing proteins (HOXs) regulate body segmentation and organogenesis during embryo development [4]. They bind to sequence specific DNA though their homeodomain [4]. In humans, there are 39 HOX genes organized into 4 clusters. Each cluster is composed of paralogous genes 1-13 [4]. The 3’ to 5’ organization of these genes reflects their temporal and spatial expression pattern during embryonic morphogenesis. The 3’ HOX genes are activated first in anterior embryonic domains followed by the expression of 5’ genes in caudal areas [4, 5]. The nested and partially overlapping expression of HOX genes in an anatomic region can determine its tissue specificity which is termed as “HOX code”[6].

HOXB13, the most posterior HOXB gene, is expressed in the caudal region of developing embryos including the tailbud, parts of spine, and hindgut as well as the urogenital sinus, from which the prostate is developed [7, 8]. In adult prostates, especially in luminal epithelial cells, HOXB13 is highly expressed [9–11]. Consistent with the general role of HOX proteins in cell fate determination and tissue differentiation[4], HOXB13 regulates prostate development and epithelial cell differentiation in normal tissues [9, 12]. HOXB13 is also involved in prostate carcinogenesis and cancer progression [13]. Studies have shown that mutations in the HOXB13 gene are associated with PCa [13]. Additionally, microarray-based transcriptome analyses revealed the progressive upregulation of HOXB13 in PCa cells and HOXB13 is considered a marker for prostatic cancer origins, to distinguish metastatic PCa from other cancers [14, 15]. However, it remains unclear whether HOXB13 plays a tumor suppressive or pro-oncogenic role in PCa. For example, HOXB13 expression is elevated in castrate-resistant PCa and induced expression of HOXB13 promotes androgen-independent growth of LNCaP cells [16]. However, constitutive expression of HOXB13 suppresses colony formation in these cells [17].

Previously, we identified genes that are differentially expressed between NEPCa and prostate adenocarcinoma (AdPCa) [18]. Among these genes is HOXB13, whose mRNA level is significantly decreased in NEPCa compared with AdPCa in all the datasets examined [18]. Here, using immunohistochemical staining, we evaluated the protein expression of HOXB13 in AdPCa and NEPCa. We discovered that HOXB13 expression is reduced or lost in NEPCa. Additionally, we present evidence supporting a change of HOX code in NEPCa. Finally, we found that the loss of HOXB13 expression in PCa cells can be reverted.

## Results

### The mRNA level of HOXB13 is decreased in NEPCa

The expression data of HOXB genes were extracted from five PCa datasets that contain both AdPCa and NEPCa samples [18–23]. As shown in Fig.1, the mRNA level of HOXB13 was lower in NEPCa samples than AdPCa in all the datasets examined. In contrast, the mRNA levels of several other members of the HOXB family including HOXB3, HOXB5, HOXB6 and HOXB7 were higher in NEPCa than AdPCa.

**Figure 1.**
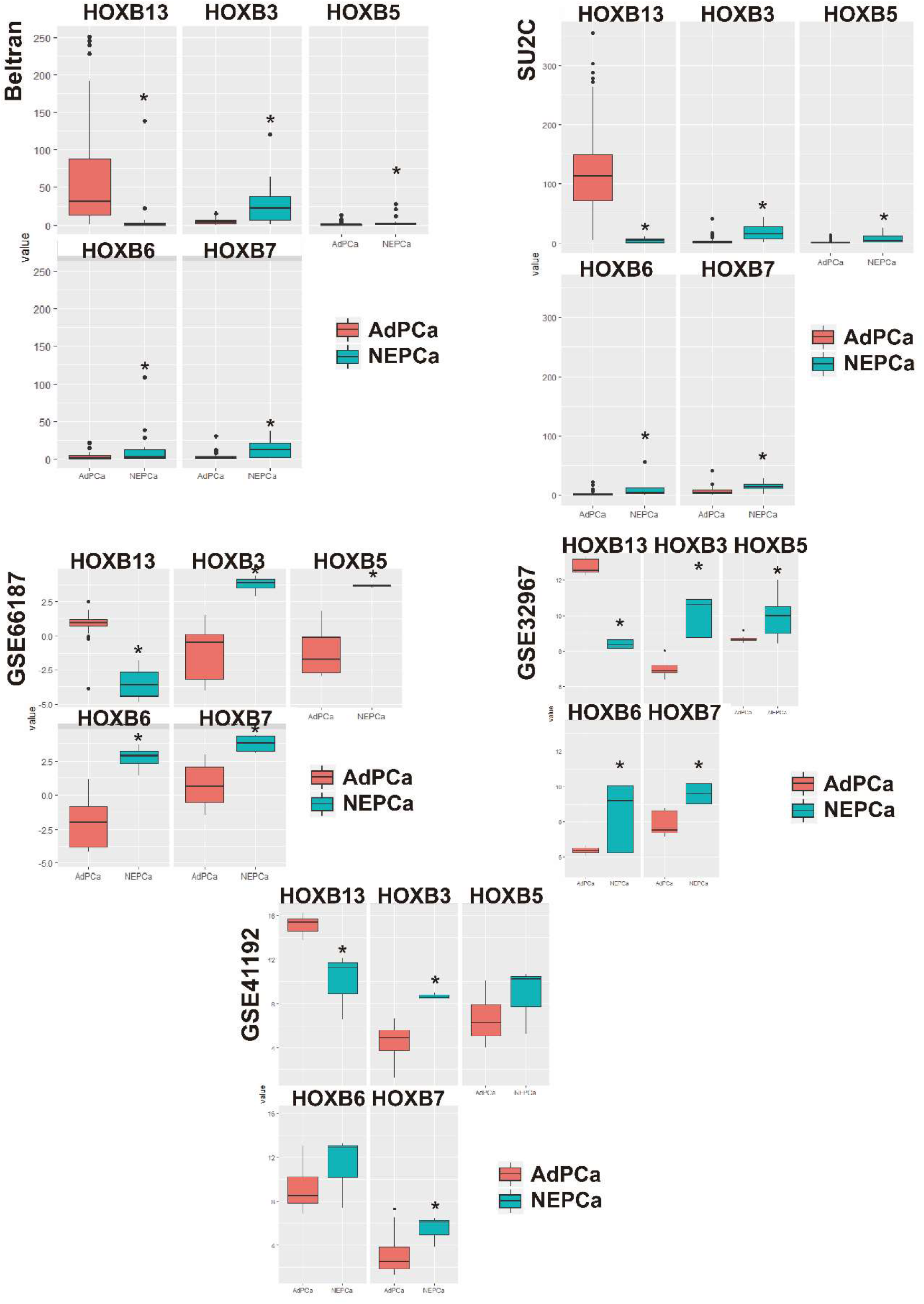
The expression of HOXB genes in AdPCa and NEPCa samples. The expression data of HOXB13, B5, B6, and B7 were extracted from NEPCa Beltran dataset [19], SU2C [20], GSE66187 [21], GSE32967 [22] and GSE41192 [23]. HOXB13 level was lower in NEPCa than AdPCa samples whereas the levels of HOXB5, B6, and B7 were higher in NEPCa samples. * p<0.05, t-test.

### The protein level of HOXB13 is lower in NEPCa

Immunohistochemical (IHC) staining was conducted to assess the protein expression of Hoxb13 in murine NEPCa tumors derived from TRAMP mice (n=10). As shown in Figs. 2A-2C, Hoxb13 expression was detected in prostatic intraepithelial neoplasia (PIN) lesions (Fig. 2A). However, the expression of Hoxb13 was decreased in the NEPCa area (Fig. 2A), which was highlighted by the positive stain of NEPCa markers Foxa2 (Fig. 2B) and chromogranin A (Fig. 2C).

**Figure 2.**
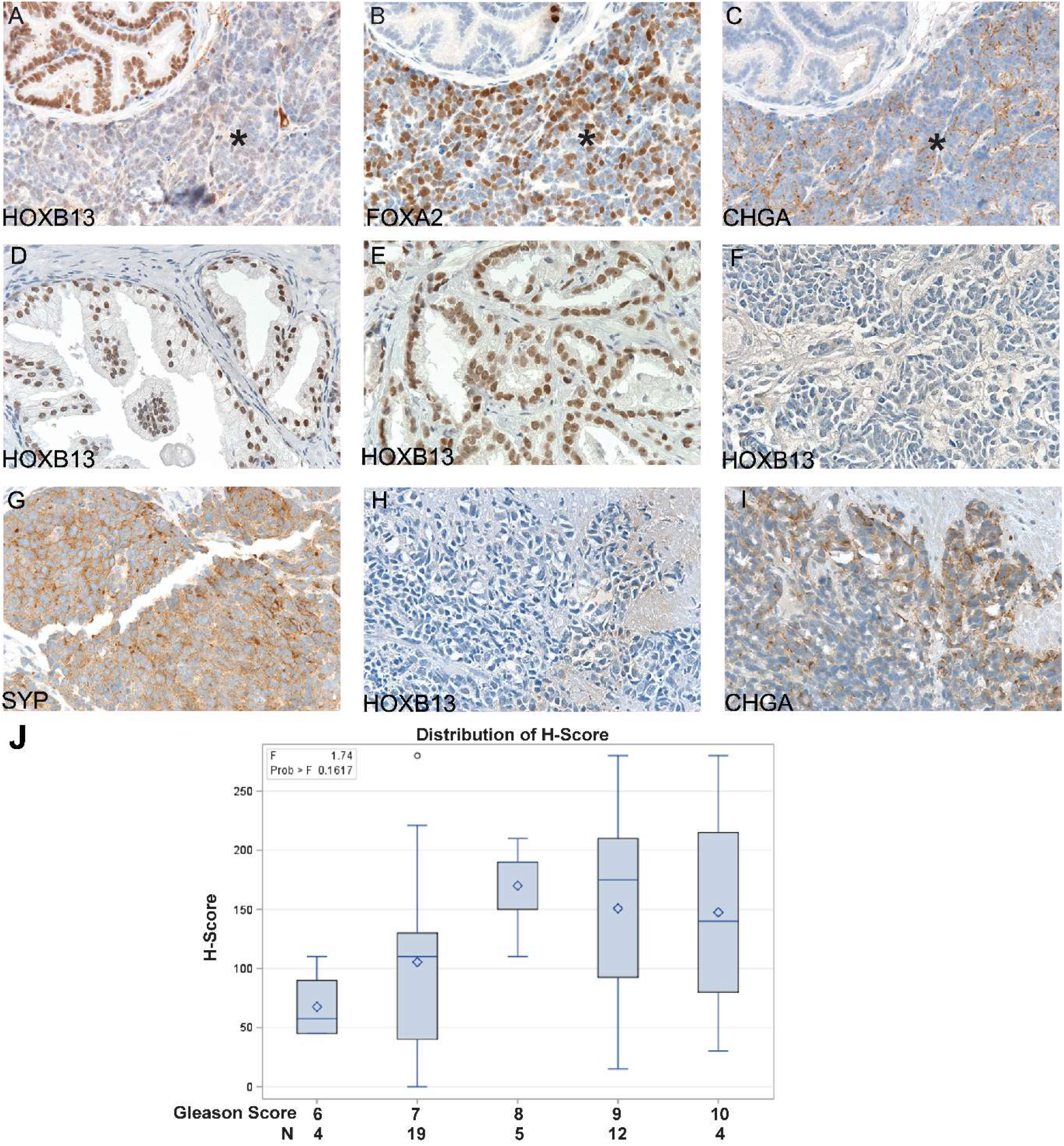
Immunohistochemical staining to evaluate the expression of HOXB13 in prostatic tissues. (A-C) Serial sections of NEPCa tumor derived from a TRAMP mouse displayed low HOXB13 expression in NEPCa area (indicated by * sign) in contrast to the positive staining in focal PIN. NEPCa markers Chromogranin A (CHGA) and FOXA2 were stained positive in NEPCa cells but negative in PIN. (D) HOXB13 expression in BPH, n=33. (E) HOXB13 expression in AdPCa, n=44. (F-I) Human NEPCa tumors demonstrated reduced (n=1) or negative (n=8) staining of HOXB13. These NEPCa tumors were stained positive for Synaptophysin (SYP) or Chromogranin A (CHGA). (J) Quantification of HOXB13 immunohistochemical staining with H-scores in prostatic acinar adenocarcinomas with a Gleason score of 6 (3+3), 7 (3+4 and 4+3), 8 (4+4), 9 (4+5 and 5+4), and 10 (5+5). The H score of HOXB13 immunohistochemical staining was correlated with Gleason score, Spearman correlation test, Rho=0.348, p<0.05.

The expression of HOXB13 was also evaluated in human prostatic tissues including benign prostate, AdPCa and NEPCa. As shown in Figs 2D-2I, HOXB13 expression was detected in both benign prostatic tissues (Fig. 2D) and prostate adenocarcinomas (AdPCa, Fig. 2E). In benign prostate, HOXB13 was stained positive in 26 of 28 BPH. Compared to that of BPH, the intensity of HOXB13 staining was greater in adenocarcinomas with Gleason scores of 7, 8, 9, and 10. However, wide variations in H-score were observed in adenocarcinomas with Gleason scores of 7, 9, and 10 (Fig. 2J). The largest variation in H-score ranged from 15 to 280 was noted in adenocarcinoma with Gleason score of 9. The highest H-score of 280 was found in adenocarcinomas with Gleason scores of 9 and 10. There was a positive correlation between the expression of HOXB13 (H-score) and Gleason scores of AdPCa (Spearman correlation test, Rho=0.348, p<0.05).

In contrast to the positive stain in AdPCa, no HOXB13 immunohistochemical reactivity was observed in 8 of 9 NEPCa including 8 small cell carcinoma and 1 neuroendocrine carcinoma, not otherwise specified. Of the 9 NEPCa, there was only 1 small cell carcinoma stained weakly and focally (H-score of 2) with HOXB13 immunomarker.

In summary, HOXB13 expression was detected in both benign prostatic tissues and prostate adenocarcinomas but its expression was lost or reduced in NEPCa tumors.

### HOXB13 is expressed in prostatic luminal epithelial cells

Dual immunofluorescence staining was conducted to examine the expression pattern of HOXB13 in prostatic glands. As shown in Fig. 3, the expressions of HOXB13 and basal epithelial cell marker (CK5 or CK14) were mutually exclusive. However, HOXB13 was co-expressed with prostate luminal epithelial cell marker NKX3-1. The luminal expression of HOXB13 was further supported by an analysis of a single cell RNA-seq dataset of normal human prostate (GSE120716), where HOXB13 positive cells were enriched in luminal epithelial subpopulation (sFig. 1) [24].

**Figure 3.**
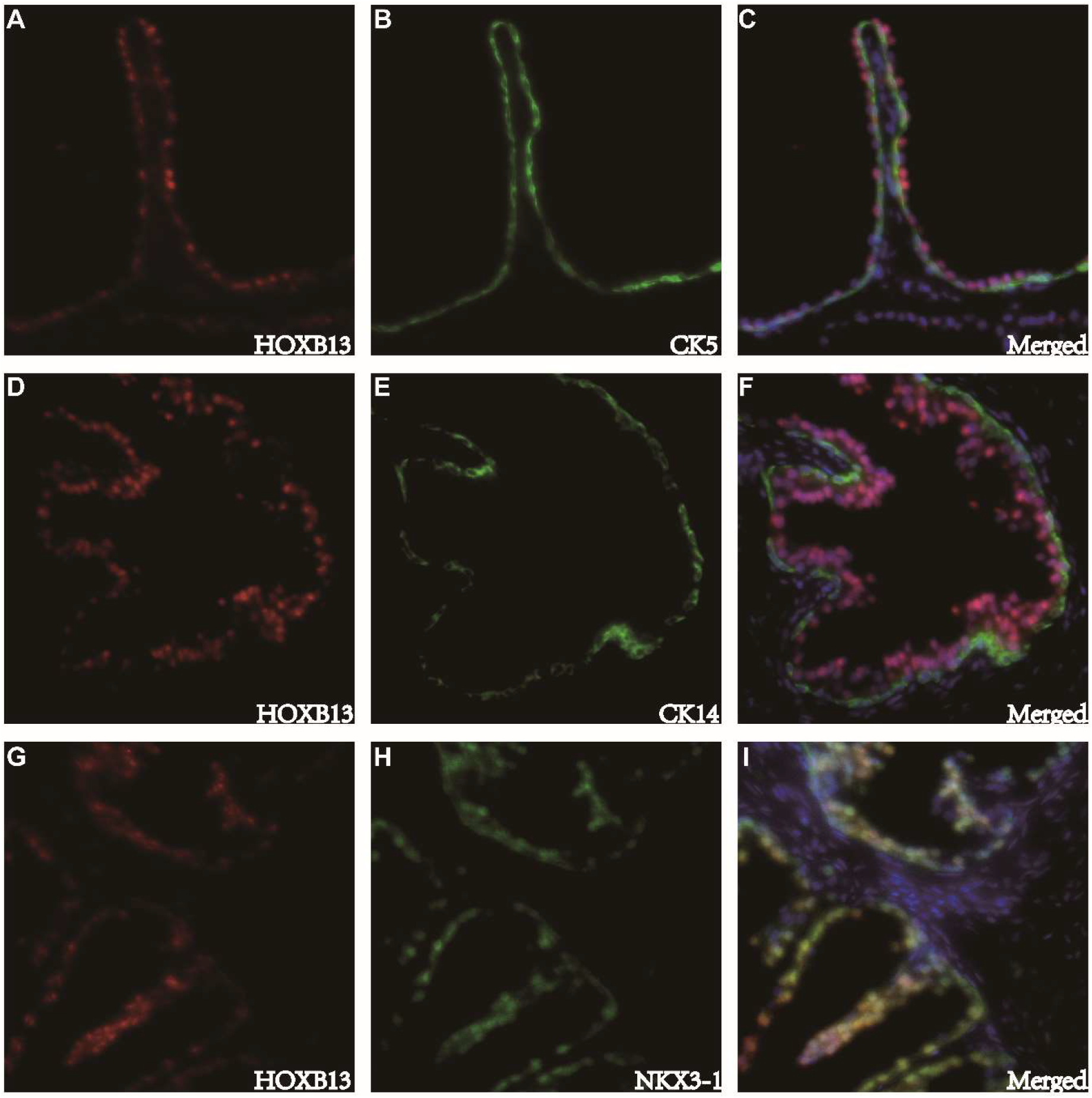
Dual immunofluorescence staining of HOXB13 with prostate basal epithelial markers cytokeratin 5 (CK5), cytokeratin 14 (CK14), or with prostate luminal epithelial markers NKX3-1. HOXB13 expression was detected in prostate luminal but not basal epithelial cells.

### Change of HOX code is associated with NEPCa

It has been suggested that different tissues have their unique HOX code. The altered expression of HOX genes in NEPCa vs AdPCa could reflect a change of HOX code in NEPCa. Using the RNA-seq data collected by the Cancer Cell Line Encyclopedia [25], we analyzed the expression of HOX genes in 1,019 established cell lines that were derived from various tissue origins and calculated the HOX code for each tissue type. The workflow is presented in Fig. 4A and the median correlation scores between cell lines of the same tissue origin and the HOX codes of various tissues are presented as a heatmap (Fig. 4B). This heatmap reflects how closely each tissue type is correlated with its HOX code. As shown in Fig. 4B, the majority of tissue types correlated well with the calculated HOX codes. Among these tissues, the most notable was the prostate group (consisting of 8 cell lines), followed by autonomic ganglia and skin, all displaying a strong correlation with their HOX code.

**Figure 4.**
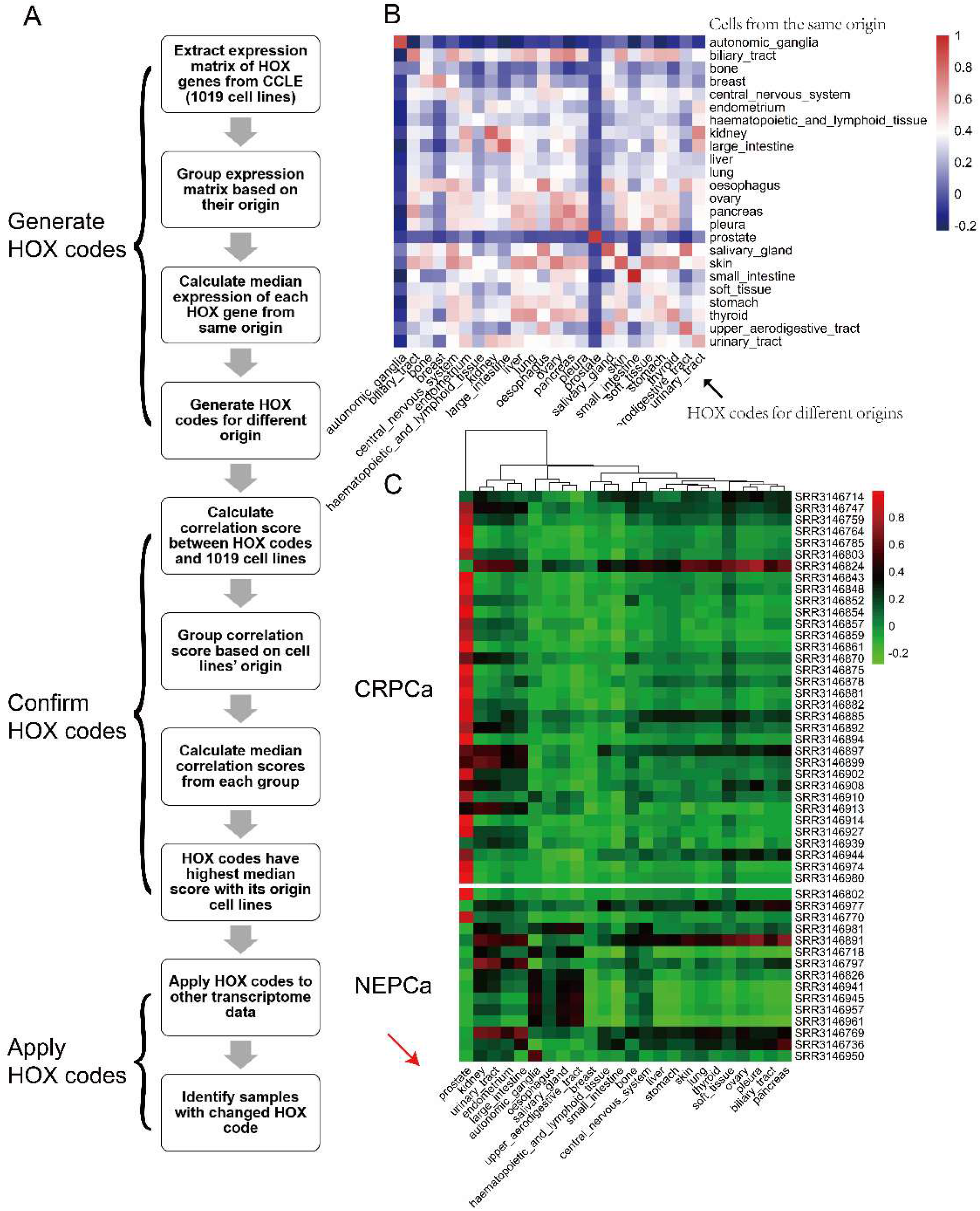
The loss of prostate HOX code in NEPCa tumors. (A) Workflow for establishing the HOX codes of different tissues. (B) The correlation of the cell lines’ tissue origins (X-axis) and their HOX codes (Y-axis). Majority tissues correlated well with their HOX codes. It is noteworthy that prostate tissue displayed a high correlation with its HOX code (C) AdPCa and NEPCa tumors displayed high and low correlation with prostate HOX code, respectively.

The strong correlation between tissue types and their HOX codes suggests that the HOX codes could be used to identify the origins of cancer tissues. Therefore, we applied the HOX codes to a RNA sequencing dataset of human PCa including 34 CRPCa and 15 NEPCa samples [2]. The correlation scores between the expression of HOX genes in each sample (row) and the HOX codes for various tissues (column) were visualized in a heatmap (Fig. 4C). The vast majority (31/34) of CRPCa samples displayed a high correlation (Pearson correlation score >0.3) with the HOX code of prostate but not of other tissues. More importantly, the majority of (13/15) NEPCa samples showed a low or negative correlation with the prostate HOX code. Similar results were also observed in other NEPCa datasets including SU2C dataset [20] (sFig 2A), GSE32967 [22] (sFig. 2B), GSE41192 [23] (sFig. 2C), GSE66187 [21] (sFig. 2D). While NEPCa tumors displayed a loss of the prostate-specific HOX code, they did not show a consistent correlation with the HOX codes of any other tissue (Fig. 4 & sFig. 2).

Taken together, these results indicate that HOX code, to certain extent, could reflect the identity of tissues and that the loss of the prostate-specific HOX code in NEPCa reflects a loss of prostatic identity in these tumors.

### DU145, a commonly used PCa cell line, is more similar to NEPCa cell line H660

Among the eight prostatic cell lines collected in the Cancer Cell Line Encyclopedia dataset [25], H660 (a NEPCa cell line), DU145 (a widely used AdPCa cell line), and PRECLH cell lines displayed a low association with the prostate HOX code whereas the other five PCa cell lines displayed a strong correlation with prostate HOX code (sFig.3). Clustering analysis of HOX genes in PCa cell lines indicates that the cell lines that have lost prostate HOX code (DU145, H660, and PRECLH) are clustered together, distinct from the other five cell lines (Fig. 5A). Compared with that of the other five cell lines, the expression levels of HOXB13, HOXC13, and HOXC6 were lower in the three cell lines that have lost prostate HOX code. The reduced HOXB13 expression in H660 and DU145 cells was confirmed by Western Blot analysis (Figs. 5B & 5C). These results indicate that DU145, labeled as an AdPCa cell line, is more similar to a NECPa cell line than other AdPCa cell lines with regards to its HOX code.

**Figure 5.**
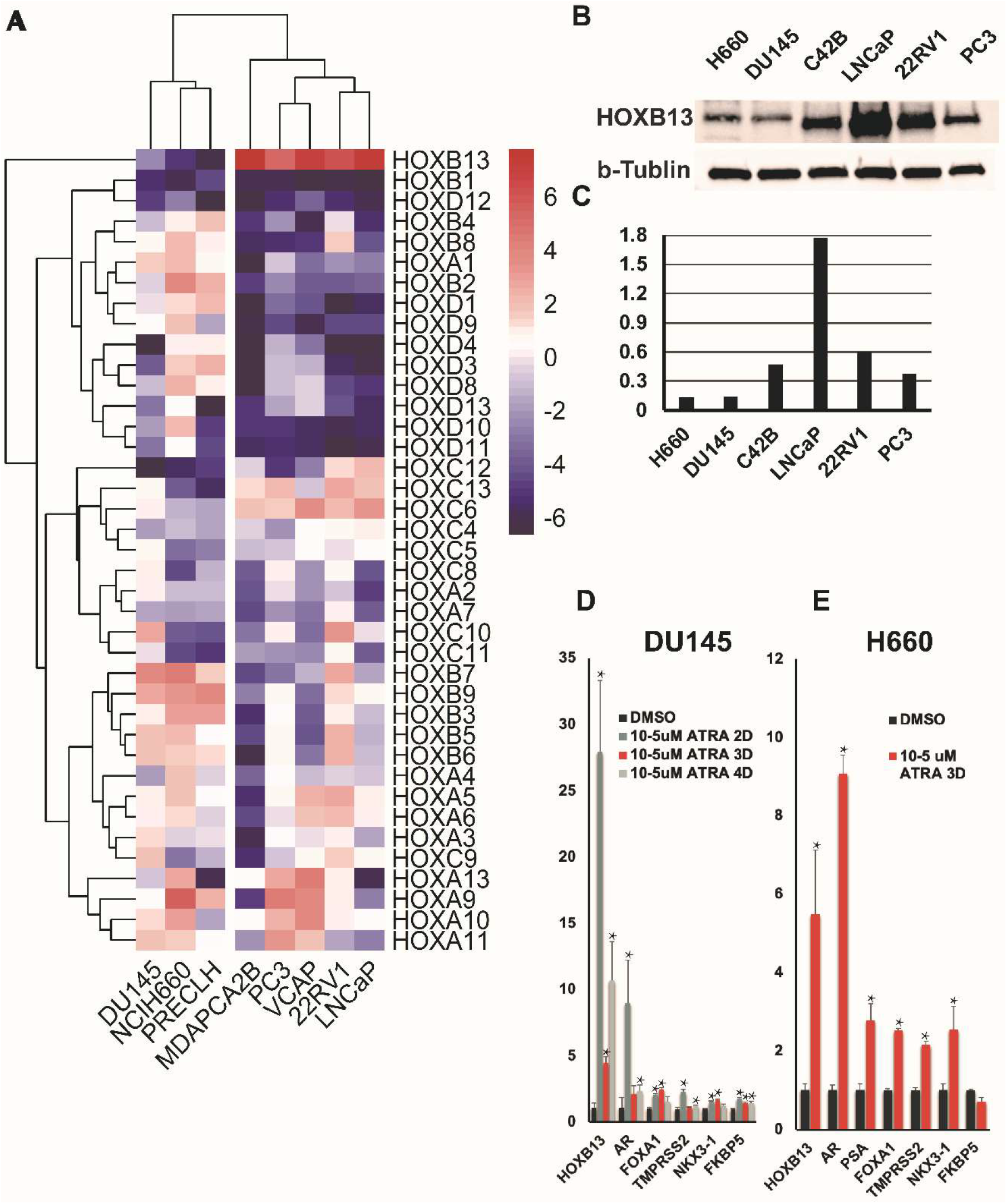
The decreased HOXB13 expression in PCa cells can be reverted. (A) A heatmap of HOX genes in PCa cell lines. DU145, H660, and PRECLH cell lines were clustered together, away from the other PCa cell lines. HOXB13 levels were lower in these three cell lines. (B) Western blot analysis confirmed the low HOXB13 protein expression in H660 and DU145 cells. (C) The quantification of Western blot result. (D & E) RT-qPCR to assess the levels of HOXB13 and AR in H660 and DU145 cells that were treated with all-trans retinoic acid (ATRA, 10^−5^ M). ATRA induced the mRNA expression of HOXB13, AR, and AR targets includingTMPRSS2, FKBP5, and Nkx3-1, but not PSA.

### The lost expression of HOXB13 in PCa cells can be reverted

Accumulating evidence supports that NEPCa tumors emerge via the trans-differentiation of AdPCa and that transcriptome reprogramming plays a key role in this process. To test whether the loss of HOXB13 expression in PCa cells results from epigenetic reprogramming and whether NEPCa cells can be reverted to AdPCa, we treated DU145 and NCIH660 cells with all-trans retinoic acid (ATRA), a commonly used agent to induce the differentiation of stem cells [26]. Here, we tested whether ATRA could induce the expression of HOXB13 as well as the expression of prostatic differentiation genes (AR and it targets) in PCa cells. The established NEPCa NCIH660 cell line was used in this experiment. Because the DU145 cell line was shown to display a similar HOX code to H660 and previous studies have shown that DU145 cells have some NEPCa features [27], this cell line was also included in the ATRA experiments.

As shown in Fig. 4D, the HOXB13 mRNA expression was induced in both H660 and DU145 cells that were treated with ATRA. An important feature of NEPCa is the loss of AR signaling; therefore, we examined the expression levels of AR and its downstream target genes in these cells. ARTA induced the mRNA expression of AR and its targets including TMPRSS2, FKBP5, NKX3-1 but not PSA in both DU145 and H660 cells (Fig. 4E). These data indicate that the lost HOXB13 expression and AR signaling can be reverted in H660 and DU145 cells.

## Discussion

In this paper, we report that HOXB13 is expressed in benign prostatic tissues and prostate adenocarcinomas, but its expression is decreased or lost in both human and mouse NEPCa. These data indicate that the loss of HOXB13 expression is a common feature of NEPCa.

HOX proteins are important transcription factors that regulate the anterior-posterior patterning in embryos [4]. A change of HOX expression results in deficiencies in the axial formation and the loss of tissue identity. In this study, using transcriptomic data of 1019 cell lines, we established a method to calculate the HOX codes for 24 tissues and applied the HOX codes to PCa RNAseq datasets. Using this method, we identified samples that uphold or lose prostate identity. We showed that prostate HOX code was sustained in the majority of AdPCa samples. This result is consistent with previous reports that HOXB13 is a PCa (adenocarcinoma) marker [14]. In contrast, the majority of NEPCa tumors and the NEPCa cell line H660 displayed low correlation with the HOX code of prostates, indicating a loss of prostate identity in NEPCa cells. DU145, a PCa cell line that has certain NEPCa features [27], also displayed a loss of HOX code. Taken together, these data suggest that during NEPCa development, the prostate identity is gradually lost in PCa cells.

Though NEPCa tumors seemed to lose prostate identity, they did not show consistent correlation with the HOX codes of any other tissues, suggesting that they have yet acquired a clear-cut new identity. This is in line with the consensus that NEPCa cells have lost prostate differentiation but this notion is not in complete agreement with the proposed acquisition of neuronal features in NEPCa. Our observation of the lack of clear-cut identity of NEPCa is an addition to this line of knowledge. Our data suggest that NEPCa cells represent a state of loss of prostate identity, but these tumor cells have not acquired or may never acquire a new identity. Although the acquired neuronal “features” in some NEPCa tumors resemble neuronal characteristics in certain aspects, NEPCa cells have not acquired the neuron “identity”.

Additionally, our HOX code method provides a unique technique to study lineage switching during cancer progression. By applying the HOX code analysis to various prostate cell lines, we found that three prostatic cell lines lose the prostate-specific HOX code. Among these, H660 is a NEPCa cell line and DU145 is a commonly used androgen-independent PCa cell line. Analysis of the HOX expression profile indicates that DU145 is more similar to NEPCa (H660) than AdPCa cell lines. Given the NEPCa features in DU145 cells, these cells may represent a unique state of PCa – they have started losing prostatic differentiation but have yet to gain a full NE phenotype. This cell line could be a good model system for inducing NE differentiation.

Currently, there are no effective treatment options against NEPCa. A possible therapeutic approach is to revert the lineage switch of NEPCa. By inducing prostatic differentiation in NEPCa cells, we might be able to revert them back to the AdPCa state and re-sensitize these cells to androgen deprivation therapy. For this purpose, we treated HOXB13-low NEPCa (NCIH660) and NEPCa-like (DU145) cells with retinoic acid, a commonly used agent to induce differentiation and a key molecule involved in body segmentation during vertebrate embryogenesis [28]. We found that retinoic acid induced the expression of HOXB13 and AR in both H660 and DU145 cell lines. This indicates the loss of HOXB13 expression during the development of NEPCa is reversible. This notion also supports the trans-differentiation theory of NEPCa. More studies would follow to determine the role of HOXB13 in the recovery of AR signaling.

Our observation on the effects of ATRA is consistent with previous reports that ATRA induces the expression of HOXB13 in DU145 cells and inhibits NEPCa in TRAMP mice, PCa mouse models [29, 30]. We showed here that ATRA reverted the lost expression of HOXB13 and AR in NEPCa cells, providing evidence supporting the possibility of inducing prostate differentiation in NEPCa cells. Further research is warranted to study the therapeutic effects of ATRA and to test the roles of other retinoic acids or differentiation-inducing agents in the treatment of NEPCa, either as single agent or in combination with androgen deprivation therapy.

## Materials and methods

### Sample collection

De-identified human prostate tissue specimens were obtained from LSU Health-Shreveport Biorepository Core, Overton Brooks VA Medical Center, Ochsner Health System Biorepository, and Tissue for Research as described previously [31]. The specimens used in this research include 33 benign prostate hyperplasia, 44 AdPCa, and 9 NEPC tissues. All the tissues were used in accordance with LSU Health-Shreveport IRB protocols. Archived tumor sections of TRAMP and 12T-10 LADY mice were used for this study.

### Immunohistochemical and immunofluorescence staining

Immunostaining was performed using Vectastain elite ABC peroxidase kit (Vector Laboratories, Burlingame, CA) as described previously [31]. Primary antibodies include HOXB13 and Chromogranin A (CHGA) (sc-28333 and sc-1488 respectively, Santa Cruz Biotechnology, Dallas, TX), Synaptophysin (SYP) (611880, BD biosciences, San Jose, CA), FOXA2 (ab 23306-100, Abcam, Cambridge, MA), cytokeratin 5 (CK5) and 14 (CK14) (904801 and 905501 respectively, BioLegend, San Diego, CA). The tissue sections were counterstained, mounted, and imaged with a Zeiss microscope (White Plains, NY). The staining was evaluated using a semiquantitative H-score. Immunofluorescence staining was imaged with a Nikon fluorescence microscope (Melville, NY).

### Bioinformatics analyses

The mRNA expression data of HOXB genes were extracted from publicly available NEPCa datasets including RNAseq datasets (Beltran’s study [19], SU2C [20], and CCLE [25]) and microarray datasets (GSE66187[21], GSE32967[22], and GSE41192 [23]). The Beltran dataset contains 49 PCa samples including 34 AdPCa and 15 NEPCa [19]. The SU2C dataset contains 113 AdPCa and 5 NEPCa samples [20]. GSE66187 contains 4 NEPCa and 20 AdPCa samples [21]. GSE32967 contains 14 NEPCa and 8 AdPCa samples [22]. GSE41192 contains 3 NEPCa and 29 AdPCa samples [23]. The expression of HOXB13 was compared in ADPCa vs NEPCa samples in these datasets. Data manning was performed by using RStudio [32, 33]. Heatmaps were generated by using “Pheatmap” package [34] and boxplots using “ggplot2” package [35].

To establish the HOX code for each tissue origin, we extracted the expression matrix (FKPM) of 39 HOX genes from CCLE dataset. This dataset contains RNA-seq data of 1019 cell lines. These cell lines are grouped base on their tissue origin information. The median expression of each HOX gene in the grouped samples was calculated to generate a HOX code for each tissue origin. We then applied the HOX codes (sTable1) to each of the 1019 cell lines by calculating the Pearson correlation scores between the expression of HOX genes in that cell line and the HOX codes of various tissue origins. A heatmap was constructed to visualize the median correlation score in each group of samples (X-axis) and their correlation with HOX codes of various tissue origins (Y-axis). We also applied the HOX codes to publicly available PCa datasets and calculated the Pearson correlation scores to reflect the correlation between the HOX expressions in each prostatic sample and the HOX codes of various tissue origins.

### Cell culture and drug treatment

DU145 and H660 cells were cultured in RPMI1640 and Prostalife medium, respectively (LifeLine Cell Technology, Oceanside, CA). All trans retinoic acid (Sigma Aldrich, St. Louis, MO) was dissolved in DMSO and diluted (1:2000) with cell culture media. The cells were treated with retinoic acid at a final concentration of 10^−6^, 10^−5^, and 10^−4^ M for 1 to 4 days.

### RNA extraction and RT-qPCR

RNA was extracted by using RNeasy Mini Kit (Germantown, MD). Reverse transcription and qPCR were conducted by using iScript™ Reverse Transcription Supermix and SYBR green PCR Supermix (BioRad, Hercules, CA). GAPDH was used to normalize the gene expression. Primer sequences were listed in sTable 3.

### Western blot

Cells were collected in PBS and lysed in passive lysis buffer (Promega, Madison, WI). Equal amounts of protein were loaded for Western blot analyses. ProSignal® Dura ECL Reagent (Genesee Scientific, San Diego, CA) and Chemidoc (Bio-Rad, Hercules, CA) were used to visualize the proteins. Beta-tubulin was used as loading control.

### Statistical analyses

T-test was used to compare the gene expression in AdPCa and NEPCa samples. Spearman Correlation Coefficient test was used to analyze the correlation between tumor grade and protein expression (H-score). p<0.05 was considered statistically significant.

## Supporting information

sFig1

sFig2

sFig3

sTable1

sTable2

## Acknowledgement

This research was supported by NIH R01 CA226285, U54 GM104940, and LSUHSC FWCC and Office of Research funding to Yu, X.

sFigure 1 Analysis of single cell RNA sequencing data (GSE120716[24]) indicated that HOXB13 was enriched in luminal epithelia (LE) compared to basal epithelia (BE), t-test, p<0.05. (Left) Heatmap to show the different cell populations that were selected based on the expression of maker genes. Red in heatmap represents read higher than 200. (Right) Boxplot to show the distribution of HOXB13 positive cells in luminal and basal cell populations.

sFigure 2

Heatmaps to show the correlation (Pearson association) of PCa samples with the HOX codes of different tissue origins. Each role represents a PCa sample and each column represents a tissue HOX code. The NEPCa samples are labeled with “*”. The prostate HOX codes are highlighted.

sFigure 3

Heatmap to show the correlation (Pearson association) of PCa cell lines with the HOX codes of different origin. NCI-H660 is a NEPCa cell line and the rest cell lines are AdPCa.

sTable 1

HOX codes for different tissue origins

sTable 2

qPCR primer list

## Notes

### Competing Interest Statement

The authors have declared no competing interest.

