## Supplementary figures and images for "Loss of HOXB13 expression in neuroendocrine prostate cancer"

### sFig1

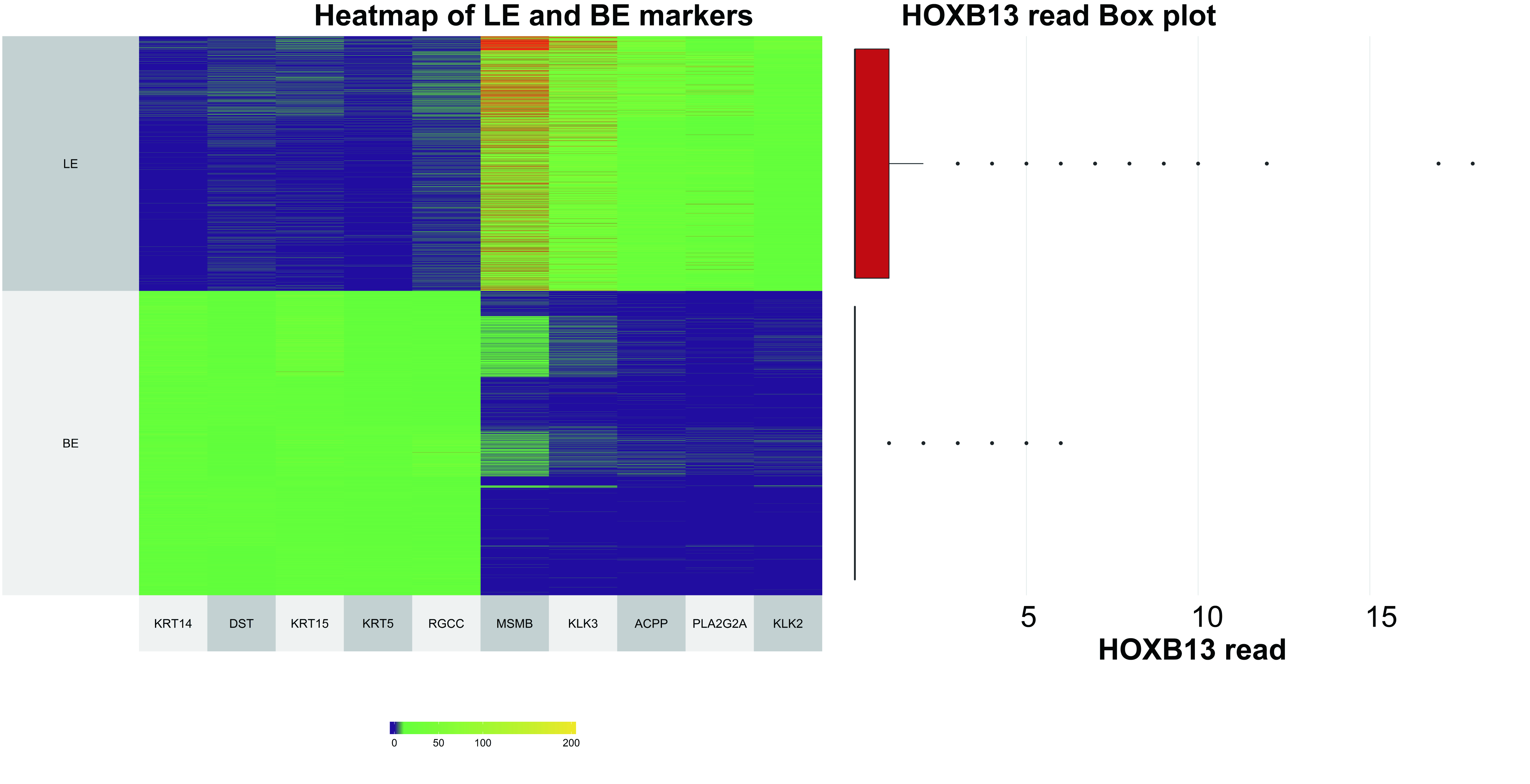

### sFig2

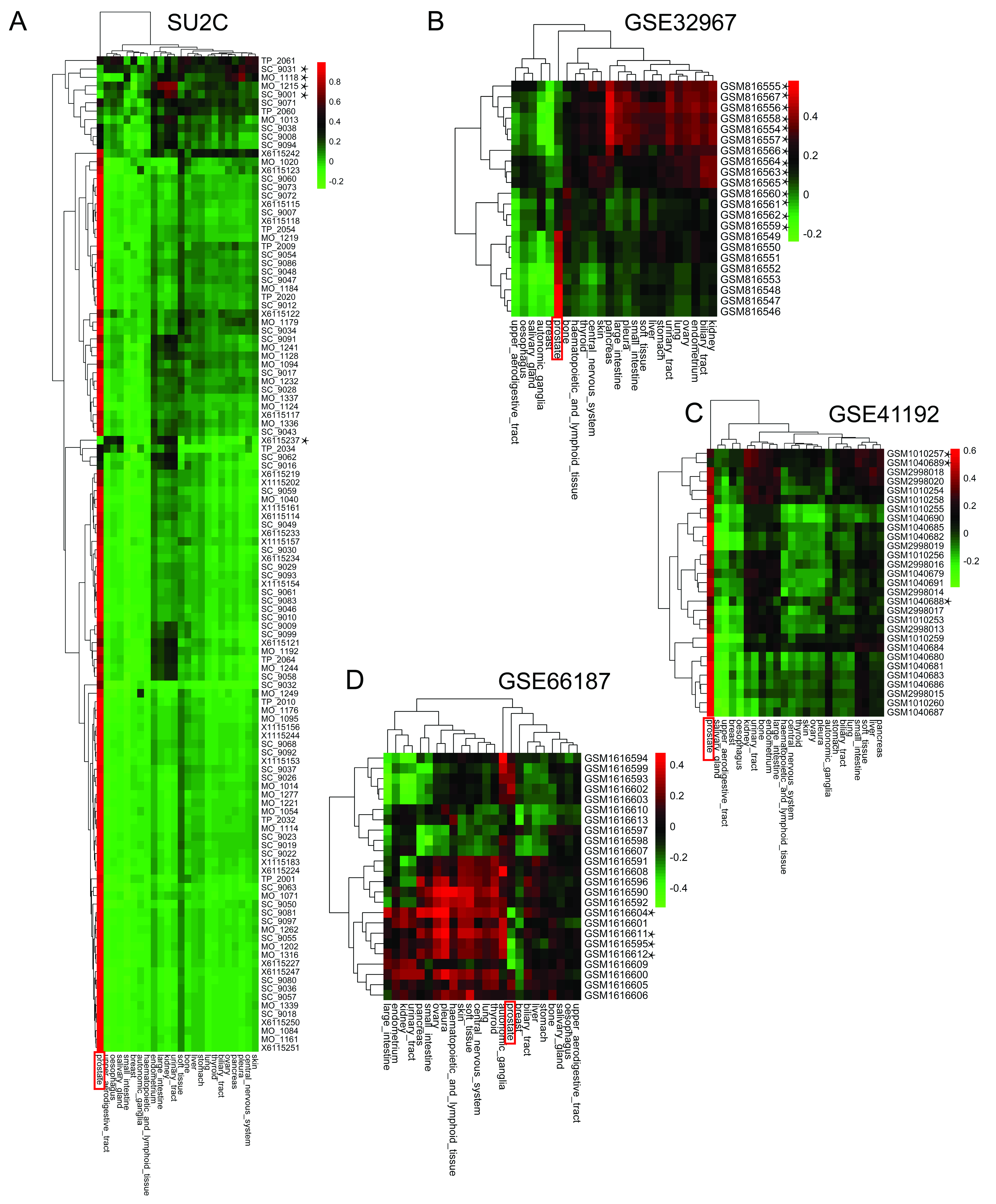

### sFig3

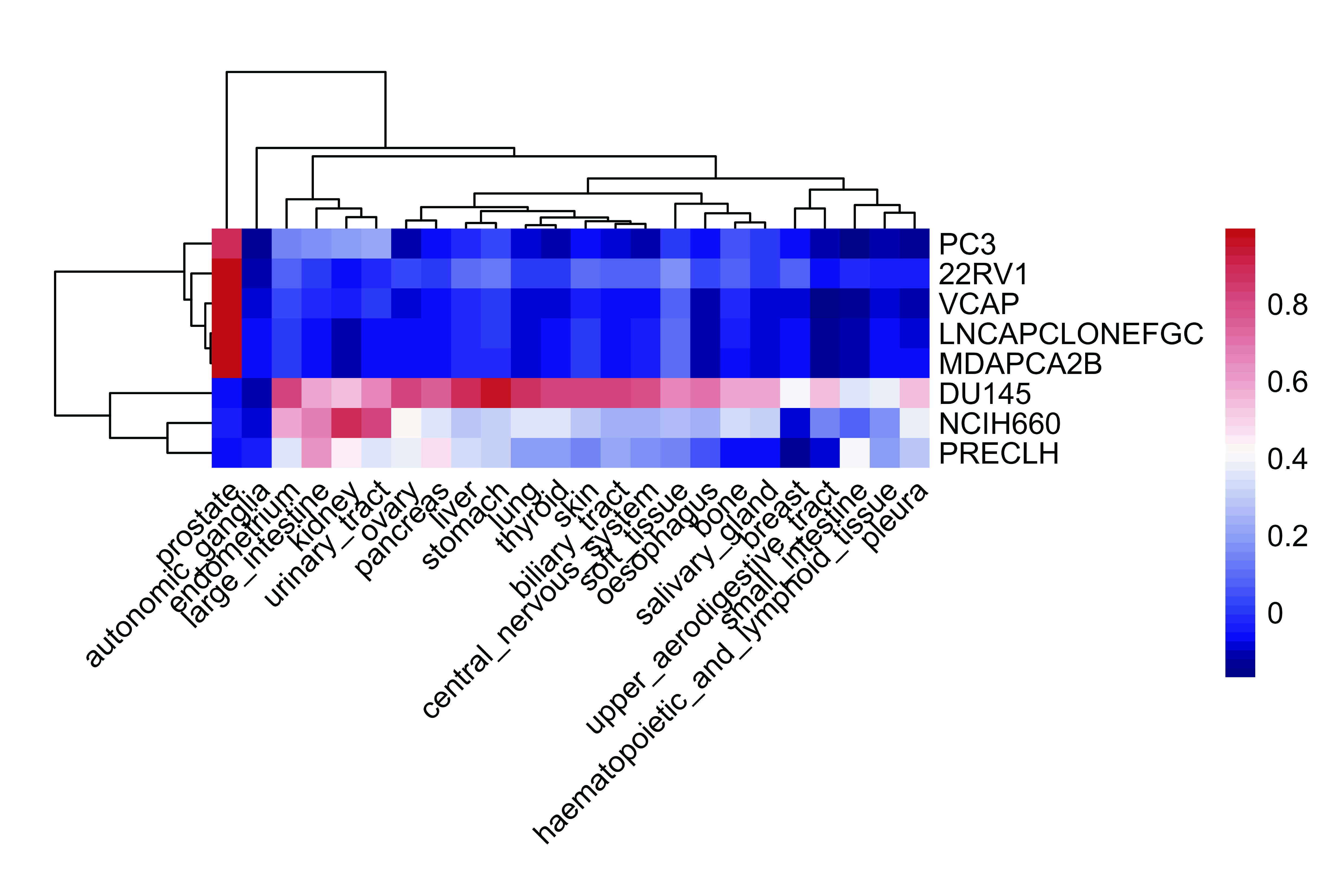
